# Identification by cryoEM of a densovirus causing mass mortality in mass-reared larval darkling beetles (*Zophobas morio*)

**DOI:** 10.1101/2022.05.14.491968

**Authors:** Judit J Penzes, Jason T Kaelber

## Abstract

A reared colony of larval superworms (*Zophobas morio*) experienced the swift and unexplained death of about 90% of its population. We isolated a high-abundance virus from dead larvae and, using cryoEM, identified it as a novel densovirus, which we name Zophobas morio black wasting virus (ZmBWV). Densoviruses (DVs) are small, ssDNA viruses of the family *Parvoviridae* that infect protostome and deuterostome invertebrates. The vast majority of DVs have been associated with severe pathology, especially in larval stage insects. By cryoEM we resolved the high-resolution capsid structure of this new DV at 2.9 Å resolution for the genome-packaging, infectious particles and at a resolution of 3.3 Å in case of the empty particles, both purified directly from the infected *Z. morio* larvae. The capsid structure of ZmBWV provides the first insights into the capsid morphology of a structurally previously-uncharacterized genus, *Blattambidensovirus*. Consequently, the ZmBWV capsid harbors a unique surface morphology within the family, yet shows the *T*=1 icosahedral symmetry, the eight-stranded jelly roll core, as well as general features of multimer interactions previously found typical of subfamily *Densovirinae*. Although we have not inoculated healthy larvae with ZmBWV, on the basis of its prodigious abundance in infected larvae and the prior probability of larval pathology with viruses of this subfamily, ZmBWV is the most probable cause of the observed mass mortality event.

## Introductionf

Parvoviruses (PV) are small icosahedral viruses with a non-enveloped virion and linear, ssDNA genome [1]. The family *Parvoviridae* possesses a remarkably diverse host spectrum of both protostome and deuterostome invertebrates as well as vertebrates. Although two out of the three subfamilies within the *Parvoviridae*—*Densovirinae* and *Hamaparvovirinae*—include viruses of invertebrate hosts, traditionally all invertebrate-infecting PVs are referred to as densoviruses (DVs). DVs, in general, display high virulence and pathogenicity, which has been shown to affect insects in their larval stage [2-8], crustaceans [9-14], mollusks [15, 16] as well as echinoderms [17-19]. Mass-reared arthropods are especially in danger of DV infection [20-24]. The superworm (*Zophobas morio*), a species of darkling beetle, is one such arthropod species; its larvae are a staple item on the diet of captive reptiles, birds and amphibians worldwide, and it is under investigation as a next-generation animal protein source [25].

Members of the family *Parvoviridae* are united by their genome size and organization and presence of certain conserved protein domains, as well as their capsid protein structure [1, 26]. All parvoviruses thus far harbor a small, single-stranded DNA genome of 3.7 to 6.3 kb in length, flanked by partially double-stranded, hairpin-like DNA secondary structures, essential for parvovirus replication. The coding region of the genome contains two major expression cassettes. The cassette closer to the 5’ end, named *rep*, is capable of expressing a varied number of non-structural (NS) proteins. The cassette closer to the 3’ end, named *cap*, encodes for one to four structural proteins (VP). The VPs share an overlapping C-terminal region and differ from each other in N-terminal extensions. Most members of the vertebrate-infecting *Parvovirinae* and all members of the *Densovirinae* harbor a phospholipase A2 (PLA2) enzymatic domain in the unique N-terminal extension (VP1u) of their largest minor capsid proteins [27, 28]. The PLA2 domain is essential for endolysosomal egress during PV and DV intracellular trafficking.

Though over a hundred high resolution structures have been determined in case of the vertebrate-infecting *Parvovirinae* [29], only five DVs have been structurally characterized. Of these, three belong to members of the subfamily *Densovirinae*: Galleria mellonella DV (GmDV) of genus *Protoambidensovirus* at 3.7 Å resolution [30], Acheta domesticus DV (AdDV) of genus *Scindoambidensovirus* at 3.5 Å resolution [31] and Bombyx mori DV of genus *Iteradensovirus* at the resolution 3.1 Å [32], all of which are type species for their respective genera. Although the VP protein sequences are highly divergent among members of the three subfamilies and lack sequence-based evidence of homologous origin [26], they do share an unambiguous jelly roll fold [33], comprising eight antiparallel β-strands linked together by loops and helices of varied length. The *T*=1 icosahedral symmetric capsid assembles form 60 VPs, in which the aforementioned loops shape the highly-variable surface morphology of PVs and DVs, even within genera [29]. At the fivefold symmetry axis, all PV capsids harbor a pore-like opening, which continues in a channel, linking the capsid lumen with the environment [29]. In case of members of the *Parvovirinae* this has been shown to be the sight of genome packaging and VP1u externalization, so that the PLA2 can carry out its function [34-36].

As a diagnostic tool, electron microscopy continues to play a role alongside sequence-based methods because of its speed, lack of reliance on reference databases, and lack of bias towards linear, circular, single-stranded, double-stranded, DNA, or RNA genomes. For example, SARS-CoV was first identified by thin-section transmission electron microscopy [37]. We used cryo-electron microscopy (cryoEM) to identify a DV of genus *Blattambidensovirus* in connection with an outbreak of mass-mortality in captive *Z. morio* larvae, designated Zophobas morio black wasting virus (ZmBWV). We have determined the near-atomic 3D capsid structure of this virus, harboring a surface morphology that is unique within the *Densovirinae* subfamily.

## Results and Discussion

### A densovirus associated with mass mortality in Z. morio larvae

At a small-sized insect rearing facility in the western United States, *Z. morio* larvae approximately two months of age and about 25 mm in length were observed to show signs of distressed locomotion, uncoordinated wiggling, and rigor followed by death. The deceased larvae quickly blackened as their inner organs lost structure, essentially becoming liquefied (Fig. 1A). Within a week from the first detection of signs, 90% of the reared larvae died in this fashion. Interestingly, an outbreak of similar pathology occurred in the *Z. morio* larvae stock of the Moscow Zoo in 2015, with PCR detection revealing a DV as the causative agent [38]. The partial genome of this DV has been deposited to the GenBank under the name Zophobas morio densovirus (ZmDV). ZmDV was first described in Hungary in 2014, with similar symptoms [39]. The DV sequence, revealed in both studies, disclosed 97% nucleotide sequence identity with Blattella germanica densovirus-like virus (BgDVLV), a member of the *Densovirinae* genus *Blattambidensovirus*. BgDVLV has a genome of over 5.1 kb in length (the length of the yet–unsequenced genome termini are unknown), and it utilizes an ambisense gene expression strategy.

**Figure 1.**
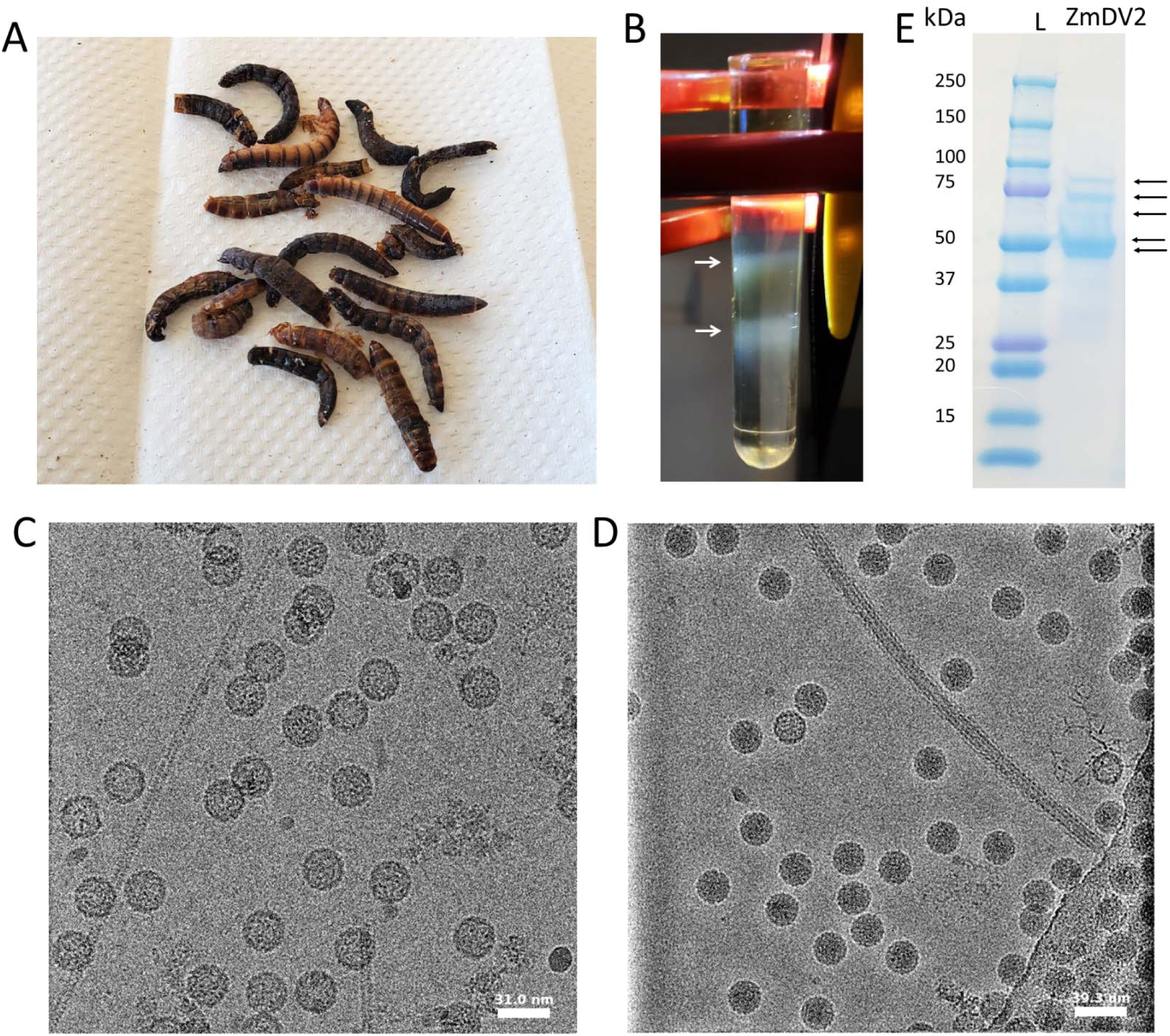
Zophobas morio black wasting virus (ZmBWV) is associated with disease in Z. morio larvae. (A) ZmBWV infected *Z. morio* larvae carcasses blacken quickly after death, with their inner organs and fat bodies quickly breaking down. (B) Virus purification from the *Z. morio* larvae from panel A, using a 5-60% sucrose step gradient, results in two well-defined bands due to particle accumulation of different buoyancy at the 20-25% (upper) and 35-40% (lower) interfaces (marked by arrows). (C) Transmission electron microscopy (TEM) micrograph showing empty ZmBWV particles from the upper band of the step gradient in panel B. (D) TEM micrograph showing genome packaging, i.e., full particles acqui ed from the lower fraction of the sucrose in step gradient in panel B. (E) SDS-PAGE of heat-denatured protein, stai ed with Coomassie brilliant blue, of the full fraction of purified ZmDV particles. The five putative capsid proteins are marked by arrows.

The *Z. morio* larvae carcasses were subjected to sucrose cushion and sucrose step gradient purification. In the step gradient, which included fractions of 5-60% sucrose at a 5% step interval, two well-defined protein bands could be observed at the 20-25% and the 35-40% interfaces, respectively (Fig. 1B). Both bands were collected, dialyzed, plunge-frozen on an electron microscopy grid. Isometric virus particles of approximately 26 nm in diameter were observed by cryoEM. This shape and diameter is consistent with our suspicion of DV infection (Fig. 1C and D). This putative DV was designated as ZmBWV. Genomic material was observed in particles from the lower buoyancy fraction but not the upper, more-buoyant fraction; the presence and buoyant separation of full and empty particles is not uncommon in non-enveloped viruses (e.g., [40]). Although ZmBWV predominated in both bands, the occasional flexivirus-shaped particle was observed. These are probably bacteriophages infecting the *Z. morio* gut microbiome.

Upon subjecting the purified particles to SDS-PAGE, five bands corresponding to the approximate sizes of 85 kDa, 74 kDa, 65 kDa, 50 kDa and 48 kDa, could be observed (Fig. 1E). Members of subfamily *Densovirinae* are known to express multiple structural proteins, with four VPs being the most common [4, 5, 20, 31, 41-43]. From VP1 to VP4, respectively, these are incorporated into the GmDV capsid in a ratio of 1:9:9:41 [30] and in the ratio of 1:11:18:30 into the AdDV capsid [31]. Based on the VP transcripts of the type member of genus *Blattambidensovirus*, Blattella germanica densovirus (BgDV), the predicted masses of three potential VPs should correspond to 85.3, 69.7 and 56.3 kDa, yet they were shown to run significantly larger due to ubiquitination. Due to leaky scanning of VP transcript 2 and 3, BgDV expresses five VPs, with VP4 being the most abundant [41]. The number and size of the ZmBWV VPs corroborates with this, further confirming that ZmBWV is indeed a blattambidensovirus. However, ZmBWV appears to incorporate an approximately equal amount VP4 and VP5 major structural proteins into its capsid and possibly lacks the aforementioned posttranslational modifications characterized for BgDV.

### The ZmBWV capsid has a densovirus-like structure with a unique surface morphology

The plunge-frozen grids with the viral particles were subjected to high-throughput cryoEM data collection to determine the atomic structure of the ZmBWV capsid by single-particle reconstruction [44]. We resolved the ZmBWV capsid structure for both the genome packaging (full), particles as well as for the empty, high buoyancy particles, at the nominal resolutions of 2.9 Å and 3.3 Å, respectively (Fig. 2A). Both structures demonstrate *T*=1 icosahedral symmetry, consistent with their family affiliation and are 28 nm at their largest diameter. Overall, the surface of the ZmBWV capsid is rather smooth, with short, yet broad protrusions surrounding the fivefold symmetry axes. Both the two- and the three-fold symmetry axes are covered with depressions. The lumen of the full particles is completely filled with electron-dense material, corresponding to the viral genome. The empty particles harbored a disordered piece of basket-like density under the fivefold symmetry axes, protruding towards the capsid lumen (Fig. 2A). Such densities have been observed previously in several parvoviral structures, including genera *Dependoparvovirus* [45-48] and *Bocaparvovirus* [49, 50] of the *Parvovirinae* subfamily and the divergent Penaeus monodon metallodensovirus (PmMDV) [51]. The basket-like density of the empty ZmBWV particles is rather similar in morphology to those found in adeno-associated virus structures, the presence and absence of which is closely-associated with the movement of the disordered VP N-termini, as a consequence of their externalization from the particle lumen [45, 46, 48].

**Figure 2.**
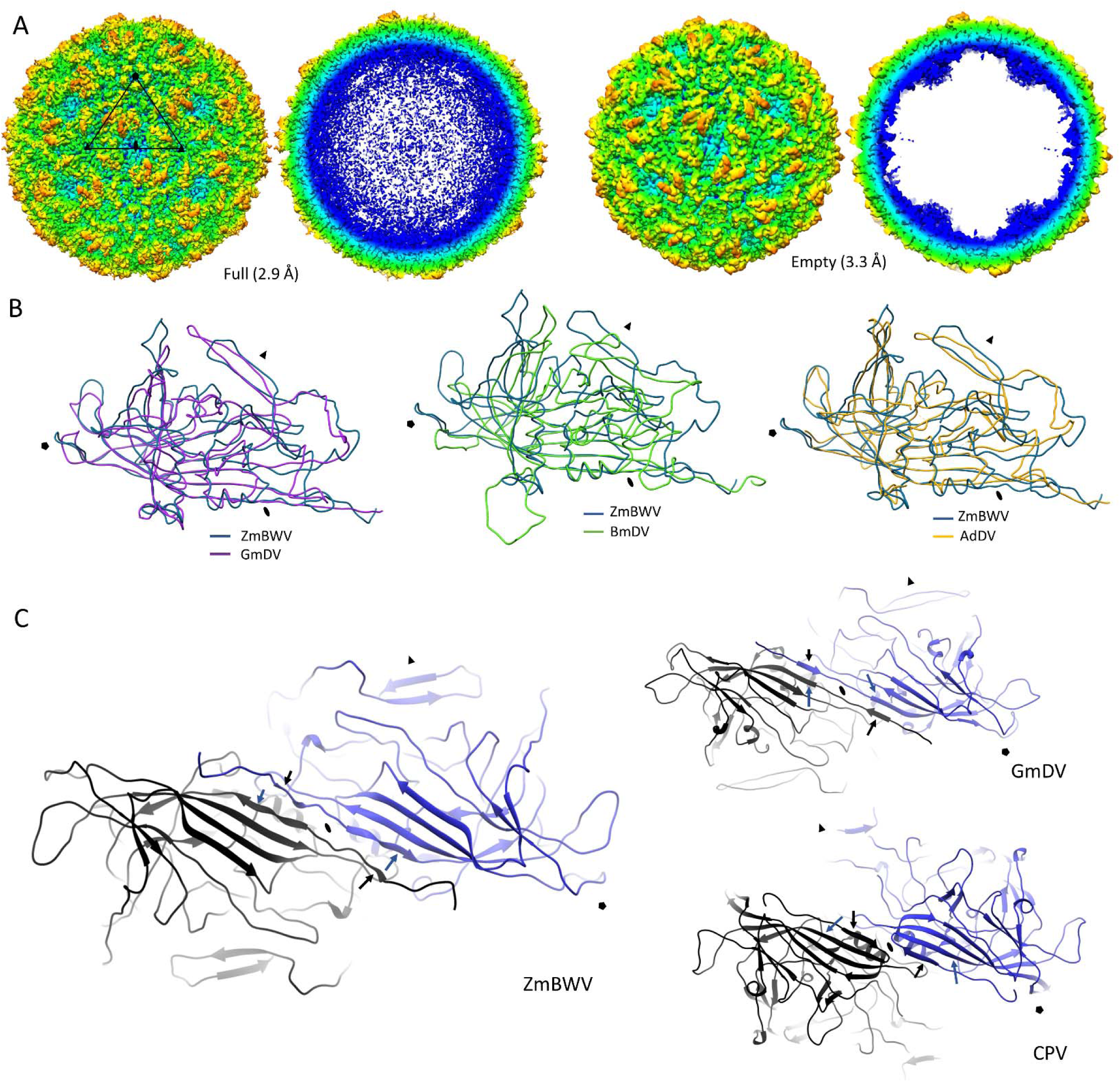
Structural studies of the Zophobas morio black wasting virus (ZmBWV) capsid. (A) Surface representation (left) and cross section (right) view of the 3D structure of the full and empty ZmBWV virus particle. The triangle represents an asymmetric unit, with a black pentagon marking the fivefold, the oval the twofold, and the two triangles the threefold symmetry axes, respectively. The map representing the empty particles is shown in the same orientation. (B) Ribbon diagram superimposition of the ZmBWV monomer backbone model with those of Galleria mellonella densovirus (GmDV), Bombyx mori densovirus (BmDV) and Acheta domestica densovirus (AdDV). Symmetry axes are labelled as in panel A. (C) Ribbon diagrams representing a dimer for ZmBWV to be compared with those of GmDV2 (*Densovirinae*) and canine parvovirus (CPV) (*Parvovirinae*). The black arrow points to the βA and the blue arrow to the βB strand in order to present the domain swapped vs. non-domain swapped dimer interactions, a fundamental difference between densoviral and parvoviral multimer interactions.

To characterize the monomeric structure of the ZmBWV capsid, we modelled the Cα backbone of the ordered protein density, which encompassed 423 residues. Inferring amino acid based on apparent side chain density, we estimate that the VP of ZmDWV is roughly 90% identical to that of BgDVLV and 60% identical to that of BgDV. The first ordered residue is located under the fivefold axis and implies the presence of a long, disordered N-terminus, in concordance with all structures of the *Parvoviridae* thus far [29]. The last ordered residue terminates at the capsid surface, in the proximity of the threefold axis, similarly to its counterparts in GmDV, AdDV and BmDV [30-32]. The structure of the monomers displayed the typical eight-stranded jelly roll fold as the core of the subunit, with surface loops corresponding with those of all determined *Densovirinae* DV structures thus far (Fig. 2B). Based on the heuristic structural comparison with all protein structures present in the RCSB Protein Data Bank (PDB), the ZmBWV monomeric structure is the most similar to that of GmDV, with a DALI Z-score of 30, followed by the AdDV and BmDV VP structures (Z-score of 29.3 and 22.6, respectively). It was equally dissimilar from those of Penaeus stylirostris DV (*Hamaparvovirinae*) and PmMDV, as well as from VP structures of the *Parvovirinae* members (Z-scores of ∼10). When the ZmBWV monomer backbone structure is superimposed with those of GmDV, AdDV and BmDV, it is apparent that the capsid surface-associated morphological differences among these four viruses are a result of various surface loop conformations, while the core itself superimposes almost perfectly (Fig. 2B).

The eight-stranded jelly roll core in most members of the *Parvoviridae* is complemented by an N-terminally located ninth strand, βA, which interacts with the luminal β-sheet, composed of the B, I, D, and G strands. Based on how this interaction is carried out, parvoviruses can adopt one of two major strategies in the way they assemble VP dimers [29]. One of these is typical of the *Parvovirinae*, in which case the βA folds back via a hairpin-like loop and interacts with the βB of the same subunit. In subfamilies *Densovirinae* and *Hamaparvovirinae* the βA is essentially a direct extension of the βB, and lacks the hairpin loop, which results in a domain-swapped conformation. The domain-swapping enables βA to interact with the βB of the twofold neighboring subunit, establishing a strong luminal twofold axis. ZmBWV displays the domain-swapped arrangement that typifies *Densovirinae* dimer interactions (Fig. 2C), and its conformation is quite similar to that of GmDV.

When comparing the capsid surface of the ZmBWV particle to those of other members of the family, it is apparent that it fits well into the general morphology of the *Densovirinae* (Fig. 3). Despite of harboring the general features of the subfamily, such as the depression at the threefold axis and a remarkably short channel at the fivefold axis, which is surrounded by rather small spike-like protrusions, its surface morphology is still distinct. These differences are the result of major conformation differences in the EF1, EF5, and GH2 loops, responsible for composing the surface spikes, as well as in the DE and HI loops at the fivefold axis and the GH3, EF3, and EF4 loops near the twofold axis. It has been shown that swapping out residues of the GH loop between two closely-related lepidopteran DVs (GmDV and Junonia coenia DV) results in mistargeted trafficking in their respective hosts [52]. As ZmBWV is the first capsid structure resolved within genus *Blattambidensovirus*, it becomes increasingly evident that DVs possess capsids differing significantly between genera, similarly to the genus-linked variability among *Parvovirinae* capsid structures [29, 53].

**Figure 3.**
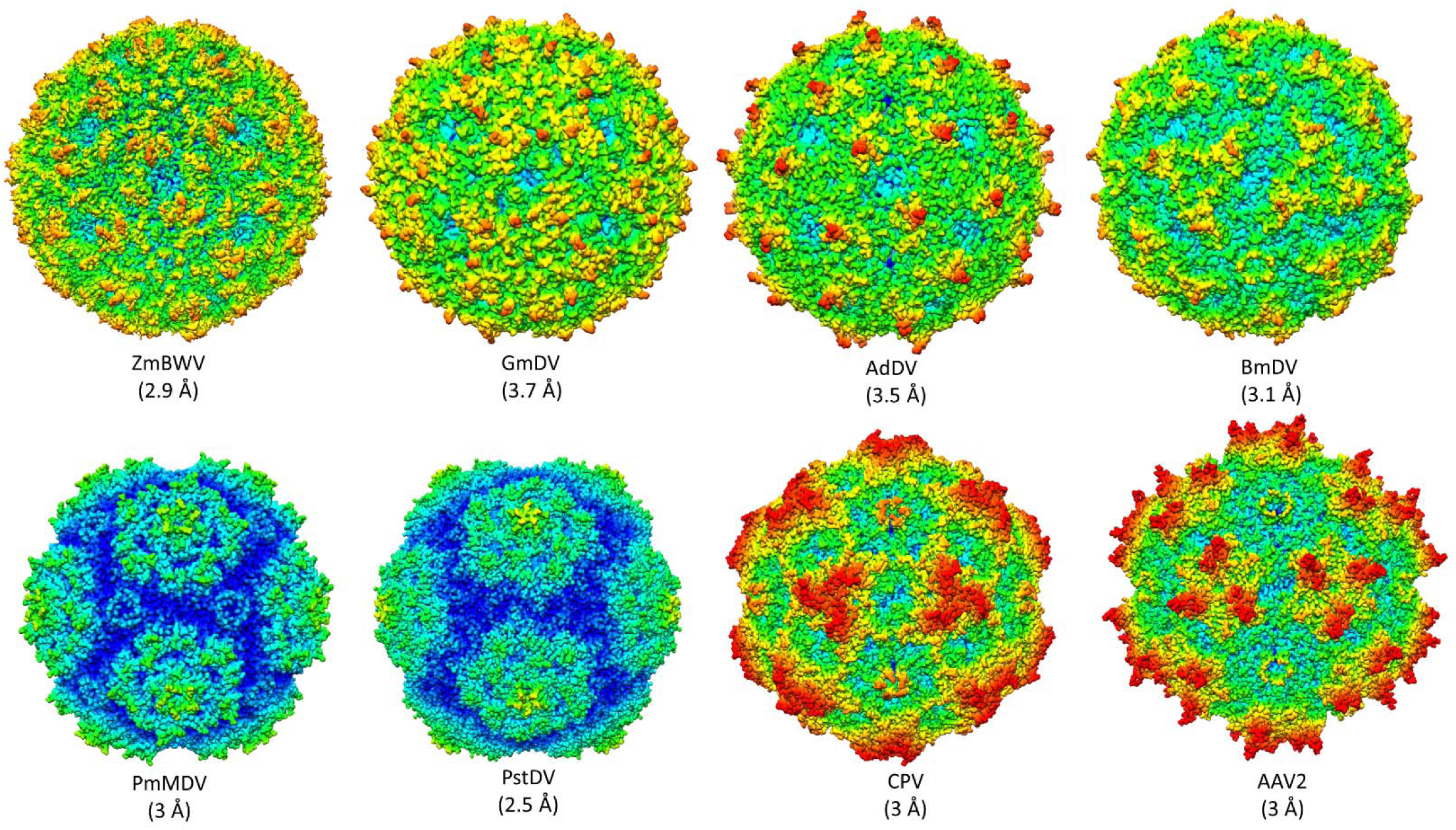
Capsid surface morphology comparison between all invertebrate-infecting, densoviral capsid structures determined thus far, along with two representatives of the vertebrate-infecting *Parvoviriane* subfamily (canine parvovirus [CPV], genus *Protoparvovirus*; and adeno-associated virus 2 [AAV2], genus *Dependoparvovirus*). Zophobas morio black wasting virus (ZmBWV) (genus *Blattambidensovirus*) described herein, as well as Galleria mellonella densovirus (GmDV) (genus *Protoambidensovirus*), Acehta domestica densovirus (AdDV) (genus *Scindoambidensovirus*) and Bombyx mori densovirus (BmDV) (genus *Iteradensovirus*) belong to subfamily *Densovirinae*, while Penaeus stylirostris densovirus (PstDV) (genus *Penstylhamaparvovirus*) is a member of subfamily *Hamaparvovirinae* and Penaeus monodon metallodensovirus (PmMDV), is of uncertain subfamily affiliation although it shares a host with PstDV. Orientation as in Figure 2A.

### Implications for husbandry

ZmBWV has not been definitely established as the causative agent of superworm liquefaction; it is possible that ZmBWV takes advantage of *Z. morio* morbidity due to some other pathogen to multiply, or that it is abundant in all populations of *Z. morio*. However, it is most probable that ZmBWV is the etiologic agent behind this disease. First, it is uncommon for a virus to grow to such high abundance in a diseased animal unless it is the cause of that animal’s disease. In this study, ZmBWV was far more abundant than all other viruses, and much more abundant than the native bacteriophages. Second, there is ample precedent for a densovirus to cause symptoms and death in farmed insects. Third, our isolation of ZmBWV from this mortality event echoes the molecular detection of ZmDV from similar outbreaks in Europe [38, 39]. Therefore, ZmBWV should be treated as the presumptive cause pending further study. We speculate that related DVs infecting *Z. morio* have a worldwide distribution.

Parvoviruses are notably resistant to alcohol-based sanitizers (unlike, for example, SARS-CoV-2) [54]. Therefore, ethanol is not sufficient for cleaning enclosures and other items that have come into contact with infected beetles. We recommend that bleach be used in these cases. Similarly, by analogy to other parvoviruses (and in contrast to SARS-CoV-2), we expect the virus to last for a long time on surfaces but not to spread particularly effectively through the air. It is likely that transmission can occur through tools and clothing as well as co-housing.

It is extremely improbable that ZmBWV poses any direct threat to humans or to pets that consume *Z. morio*. Of course, moribund larvae and liquefied larvae are unsuitable as feed for other reasons such as secondary bacterial infections.

It is unknown whether ZmBWV can be carried by asymptomatic beetles. There is some precedent for asymptomatic carriage of DVs [24]. Therefore, when bringing new beetles into a breeding colony, avoiding overtly-symptomatic individuals might not be sufficient. As a best practice, when introgressing exogenous stock into a colony, housing part of the colony separately for a couple generations is advisable to avoid loss of the whole colony in case the new beetle(s) carry this or other pathogens. Heritable immunity is conceivable but poorly-studied for this type of virus.

## Materials and Methods

### Purification of viral particles

Deceased *Z. morio* larvae were subjected to tissue homogenization in 1× phosphate-buffered saline (PBS), followed by three cycles of freeze-thaws. Following this, the homogenate was combined with an equal volume of 1× TNTM pH8 (50 mM Tris pH8, 100 mM NaCl, 0.2% Triton X-100, 2 mM MgCl_2_) and the debris were removed by centrifugation at 3700 × *g* at 4^⍰^C for 15 min intervals until the supernatant was sufficiently cleared. The supernatant was mixed with 1× TNET pH8 (50 mM Tris pH8, 100 mM NaCl, 0.2% Triton X-100, 1 mM EDTA) in a 1:3 ratio and concentrated on a cushion of 20% sucrose in TNET using a type 45 Ti rotor for 3 h at 4 °C at 42,300 rpm on a Beckman Coulter S class ultracentrifuge. The pellet was resuspended in 1 mL of 1× TNTM pH8 and, following overnight incubation, purified on a 5 to 60% sucrose step gradient for 3 h at 4 °C at 35,000 rpm, using the same instrument with a SW 41 Ti swinging bucket preparative ultracentrifuge rotor. Both visible bands were aspired by a single needle puncture and a 10-mL volume syringe. The purified fractions were dialyzed against 1x PBS in order to remove the sucrose.

### Preparation of cryoEM grids and plunge freezing

Quantifoil R1.2/1.3 300 mesh grids were glow discharged and coated with a 2.62-nm-thick carbon film. The film was fabricated by electron-beam deposition on cleaved mica using a Leica EM ACE600 instrument and floated onto a surface of ultrapure water through which the discharged grids were lifted. Samples were plunged-frozen into liquid ethane using a Vitrobot Mark IV (FEI) at 100% humidity and ambient temperature. The grids were clipped into autoloader grids and imaged using a Talos Arctica transmission electron microscope (TEM) (Thermo Fisher), equipped with a Gatan K2 direct electron detector, operated in low dose mode.

### Collection of high-resolution data and 3D reconstruction

The selected cryoEM grids were subjected to high resolution data collection, using the aforementioned electron microscope operated at 200 kV, with a 10-s-long exposure and a total dose of 43.16 e-/Å^2^, using a frame length 0.2 s. Movie frames were recorded in counting mode using the Serial EM suite [55] at a sampling of 1.038 Å/pixel. The collected movies were aligned by the MotionCorr2 application with dose-weighting [56]. The cisTEM software was used for single-particle image reconstruction [57]. Micrograph quality was assessed by CTF estimation using a box size of 512. The subset of micrographs with the best CTF fit values were included in further processing. Particles were automatically boxed by the particle selection subroutine, at a threshold value of 2.0. Boxed particles were subjected to 2D classification, imposing icosahedral symmetry at 35 classes. Particles of classes, which failed to display a clear 2D-class average of the icosahedral particle were eliminated from the reconstruction, resulting in the incorporation of 42,219 and 2265 particles in the full and empty capsid reconstructions, respectively. *Ab initio* model generation was carried out in 40 iterations, imposing icosahedral symmetry. The obtained startup volume was subjected to automatic refinement under icosahedral constraints and underwent iterations until reaching a stabile resolution. The final maps were achieved by sharpening at a post cutoff B-factor of 20. The resolution of each reconstructed map was calculated based on a Fourier shell correlation (FSC) of 0.143. The obtained cryoEM maps were visualized in Coot [58] to model the backbone of one subunit. Visualization was carried out by UCSF Chimera [59].

## Acknowledgments

We thank the perspicacious individual, who wishes to remain anonymous, who first noted these symptoms in his larvae and sent us specimens for further study. We are grateful to REGENXBIO Inc. for their support of parvovirus research in this laboratory. We thank the Institute for Quantitative Biomedicine for intramural research support funding. We thank Dr. Emre Firlar for research infrastructure support.

